# Ecological Impacts of Additive-Enriched LDPE Microplastics in Agricultural Soils: Single and Multi-Species Assessments

**DOI:** 10.64898/2026.07.17.739118

**Authors:** Blanchard Adrien, Faburé Juliette, Bamière Antoine, Breuil Sébastien, Creuzé des Châtelliers Charline, Delort Abigaïl, Etiévant Véronique, Gervaix Jonathan, Marret Manon, Plessis Cédric, Amélie A.M. Cantarel, Richaume Agnès

## Abstract

Low-density polyethylene (LDPE) microplastics (MPs) are the most frequently sampled type of microplastic in agricultural soils, potentially threatening the soil environment. The majority of MPs that have been investigated are produced from standard polymer formulations, for which the nature of the added compounds is often unknown. Furthermore, standard ecotoxicity tests performed on model species are insufficient for assessing the ecological consequences of MPs contamination in soil. This study examined the responses of multiple keystone species to exposure to MPs in interaction with various additives. No significant effects on their growth were observed when organisms were exposed to MPs alone. However, significant reductions in growth occurred when organisms interacted within uncontaminated soil: the introduction of plants reduced potworm biomass by 49 ± 4.1 % while the introduction of potworms reduced earthworm biomass by 41 ± 5.2%. In MP-contaminated soil containing plants, the average individual biomass of potworm increased significantly from 1.16 ± 0.09 mg in uncontaminated conditions to 2.01 ± 0.27 mg. This suggests that MPs limited the negative effects of interactions. Similar patterns were observed for the potworm-earthworm interaction. MPs containing the highest concentrations of additives induced the strongest biological responses. Analysis of soil parameters revealed that these impacts are likely linked to the disruption of nitrogen cycling. Therefore, it is imperative to comprehensively address the interactions between soil organisms and the influence of additives on plastic ecotoxicity in order to better assess the ecological risk posed by MPs.

## 1. Introduction

Since the 1970s, there has been an exponential increase in the production and usage of plastics, rising from 50 million metric tons to 400.3 Mt in 2022 (Geyer et al. 2017). The extensive use of plastics, coupled with suboptimal recycling efficiency and inadequate waste management, has led to the introduction of plastics to the soil environment (Barnes et al. 2009; Büks et Kaupenjohann 2020). Exposure to environmental factors such as UV radiation and temperature variation causes plastic debris to break down into smaller particles over time until they become smaller than 5 mm. At this point, the term "microplastics" (MPs) is used (Weinstein et al. 2016; Yakimets et al. 2004; Küpper et al. 2004).

Soils are now recognised as a significant reservoir of contamination. Numerous sources of microplastics have been identified in agricultural soils, including plastic mulching films (Huang et al. 2020), sewage sludge applications (Corradini et al. 2019; van den Berg et al. 2020) and recycled wastewater (Koutnik et al. 2021). Therefore, they are considered to be the main contributors to the MPs pollution in soil. A recent meta-analysis reported that the global average concentration of MPs in agricultural soils is 4.5 mgkg-1 dry soil (Büks et Kaupenjohann 2020). As demonstrated in a study by Fuller and Gautam (2016), highly contaminated sites have exhibited an average MPs concentration of 0.24% w/w. Other authors have predicted that, under a "business as usual scenario", the accumulation of plastics in soil could increase to 0.0168% and 0.1159% after 50 to 100 years from the present day, respectively (Meizoso-Regueira et al. 2024).

Low-Density Polyethylene (LDPE) is one of the most widely produced and used plastics (2019 - OECD.stat). LDPE and linear low-density polyethylene (LLDPE) account for 54,303 Mt, equivalent to 12% of the total global plastic production (see S1 for metadata of the query). As stated by Zhang et al. (2016), it is one of the most frequently sampled MP in agricultural soil. This is because it is the most widely used plastic in producing plastic mulching film and greenhouse structures. LDPE MPs have been identified as an emerging threat to terrestrial ecosystems, causing multiple adverse effects on soil biophysical properties (de Souza Machado et al. 2018; Lincmaierová et al. 2023), soil fauna (Cui et al. 2022; Rillig et al. 2019; Wei et al. 2022) and soil microorganism activities (Y. Ma et al. 2023; Rong et al. 2021). However, most studies have examined the effect on a single species, often focusing on organisms that are not ecologically significant. Overlooking other species that are responsible for essential environmental processes from which certain ecosystem services derive.

The influence of microorganisms is considered pivotal in the cycling of nutrients. As demonstrated by Wu et al. (2024), mineralisation and soil respiration are significant factors in the dynamics of soil carbon. Nitrification and denitrification are microbially driven processes that regulate the distribution, fixation and losses of nitrogen in soils (Geisseler et al. 2010). Additionally, earthworms contribute to a variety of soil functions (Kibblewhite et al. 2007; Lavelle et al. 2006; Fründ et al. 2011; Didden 1993; Blouin et al. 2013). Through their burrowing activity, they enhance soil aggregation (Blanchart et al. 1999) and influence soil carbon dynamics by interacting with microbial communities (Ingham et al. 1985; Seeber et al. 2008). Furthermore, they participate in nitrogen cycling through N mineralization (Bityutskii et al. 2002; Willems et al. 1996), yet their gut also provides an anaerobic environment for denitrifying microorganisms (Drake et Horn 2006). Earthworms are closely associated with plants through root interactions, enhancing nutrient availability and promoting stable soil structure (Scheu 2003). Numerous studies have described the positive influence of earthworms on crop yields (Baker et al. 1997). Despite the evidence showing that multispecies tests can reveal unknown indirect effects (De Laender et al. 2009; DeNoyelles et al. 1982), little research has examined the impacts of MPs on two or more interacting organisms. For instance, the presence of earthworms has been observed to mitigate the negative impact of LDPE-MPs on plant growth (Shi et al. 2024). Several studies have reported that LDPE-MPs can alter the equilibrium between soil, plants and microorganisms. Ranauda et al. hypothesized that increases of mineral nitrogen (nitrites and nitrates) after LDPE addition to soil, has facilitated the growth and activity of microbial communities involved in nitrogen cycling (nitrification and denitrification)(Ranauda, Zuzolo, et al. 2024). Therefore, LDPE MPs likely enhance all N biogeochemical cycle-related functions (Ranauda, Tartaglia, et al. 2024; Rong et al. 2021).

In this study, MPs are composed of a primary polymer such as low-density polyethylene (LDPE) mixed with a large group of compounds known as additives, which are incorporated to the plastic to impart specific properties. These additives include a wide range of substances, including dyes, slip agents, plasticisers, antioxidant stabilisers, light stabilisers, lubricants, antistatic agents, and anti-ageing agents (Polymers of Low Concern – OECD, no date; OECD 2014). It is estimated that around 10,000 substances, including polymers, are used in plastic production (Wiesinger et al. 2021). Research has demonstrated that MPs additives have the capacity to leach into the environment (Haider et Karlsson 1999; Yu et al. 2024). Many additives have been hypothesised to contribute to the toxic effects of MPs exposure (Sajjad et al. 2022), and it has been demonstrated that hydrophobic additives can act as carriers for other contaminants (Kwon et al. 2017; Fajardo et al. 2022). Despite the wide variety of substances, only a few of them have been investigated, notably phthalate esters (PAEs) and Bisphenol A (Gong et al. 2021; B. Li et al. 2023). This data highlights the urgent need to understand and quantify the impact of additives used in plastic production on the ecological effects of microplastics in soil.

This study aims to: 1) assess the toxicity of two LDPE MPs formulations on ecologically relevant soil organisms when exposed individually by comparing two additive concentrations and 2) evaluate the ecological consequences of LDPE MPs contamination in a multi-species system, to enhance the understanding of their combined functional impact. Our hypothesis is that MPs will alter the interactions between soil organisms and disrupt soil functioning, despite no effects being observed on organisms exposed individually under environmentally relevant exposure scenarios.

## 2. Materials and methods

### 2.1 LDPE Microplastics

The plastics pellets were custom-produced by Armines-CEMEF and Nice Chemistry Institute, France. Two batches were produced : (i) a standard batch (PE1) containing polymerised PE and standard dose of additives (Irganox 1076 – 0.2% (w/w) ; Irgafos – 0.1% ; Chimassorb 944 – 0.5% ; Erucamide – 0.05% ; Oleamide – 0.05% ; Glycerol monostearate – 0.25%) and (ii) an “enriched” containing ten times the standard dose of additives (PE10) (Irganox 1076 – 2% ; Irgafos – 1% ; Chimassorb 944 – 5% ; Erucamide – 0.5% ; Oleamide – 0.5% ; Glycerol monostearate – 2,5%). Formulation of the batches and purpose of the used additives are reported in S2. None of these additives has been reported as an endocrine disruptor. The plastic pellets were pre-frozen in liquid nitrogen, then ground and sieved to obtain a particle size distribution between 20 and 300 µm.

### 2.2 Soil

The soil used in this study originated from the long-term field experiment QualiAgro (a partnership between INRAE and Veolia Recherche & Innovation, Feucherolles, France - 48 53’ N, 1 58’E), established in 1998(Houot et al. 2002). The soil—an agricultural sandy-loam (loess decarbonated luvisoil) collected from the 0–30 cm horizon of the control plot—has not received any organic amendment since 1998 and is exposed only to atmospheric micro-plastic deposition, so it can be regarded as essentially uncontaminated or only very lightly contaminated. Soil samples were collected on 24 January 2023 and 11 September 2023. The soil was kept moist to preserve microbial activities, then sieved to 2mm to remove rocks and plant debris. Soil was stored at 4°C until spiking. Key soil properties include a pH value of 6.6 ± 0.1, organic carbon content at 1.03 ± 0.03 % and texture is mostly represented by silt (78.15 %), followed by clay (15.15 %) and sand (6.7 %). More details on the soil physico-chemical properties are reported in S3.

### 2.3 Soil spiking and experimental systems setup

Soil moisture was determined according to ISO 11465:1993. The MP mixture was thoroughly blended with soil in a cement mixer to achieve a final concentration of 0,1% (w/w equivalent dry soil). Organisms will be exposed to this single concentration of MPs. The spiked soil was stored at 4°C for 10 days after being moistened to a relative humidity of 70%, and 1.2kg equivalent dry soil was placed into each pot (h = 22, l = 11, w = 11 cm).

### 2.4 Multi-species experiment and tested organisms

Test soils were collected from an experimental agricultural parcel. We studied the naturally occurring soil microorganisms, focusing particularly on those involved in the carbon © cycle (respiration) and nitrogen (N) cycles (nitrification and denitrification), given their crucial contribution to soil ecosystem services as soil fertility through equilibrium of mineral nitrogen compounds available for plant growth. Two endogeic annelid species one from the macrofauna (*Aporrectodea caliginosa)* and one from the mesofauna (*Enchytraeus albidus*), were also included in the experimental design because of their key roles in organic matter transformation and soil stabilization. *A. caliginosa* and *E. albidus* are common soil-dwelling species in cultivated fields (Nowak 2004; Peigné et al. 2009; Joschko et al. 2006; Bart et al. 2018). Finally, *Lolium perenne L*., a common monocotyledonous forage plant, was included in the design to assess their influence of MPs contamination on plant growth, as plants are strongly interact with earthworms (Scheu 2003) and soil microorganisms (Guyonnet et al. 2017; Malique et al. 2019) to provide multiple ecosystemic services (Faucon et al. 2017; Pereira et al. 2018).

#### 2.4.1 Plants

Ryegrass seeds (*Lolium perenne L.*) were purchased fromVILMORIN© seeds (lot LO5254 - XDK23DT0865 - EMB 26252) and germinated for 10 days prior to the experiment in a standard potting substrate. Seedlings of uniform size were selected and planted at a rate of one per pot. In this paper, *L. perenne L*. will be referred to as “plant”.

#### 2.4.2 Worms

Two species of endogeic worms were used: the earthworm *Aporrectodea caliginosa* and the potworm *Enchytraeus albidus*. *A. caliginosa*, a common soil-dwelling species in cultivated fields, was bred at the ECOSYS laboratory (INRAE-AgroParisTech, Palaiseau, France) in an uncontaminated loamy soil collected from the Grands Closeaux site (Versailles, France). A detailed description of the breeding procedures can be found in(Bart et al. 2018). *E. albidus* were provided by Aquaplante.fr. Four *A. caliginosa* individuals were weighed and an equivalent biomass equal to 2.5 times the total potworm biomass was added to each pot at the start of the experiment. In this paper, *A. caliginosa* will be referred to as “earthworms” and *E. albidus* will be referred to as “potworms”.

### 2.5 Experimental design

The selected set of organisms **(Table 1)** were exposed to spiked soil for eight weeks at 19 ± 1°C under a 16/8 h light-dark photoperiod. Pots were watered by sub-irrigation to minimise microplastic loss through surface runoff or sedimentation. Experimental design is presented in **Fig. 1**. At the end of the experiment, soil samples were collected and stored at 4°C in order to assess microbial activities.

**Fig 1.**
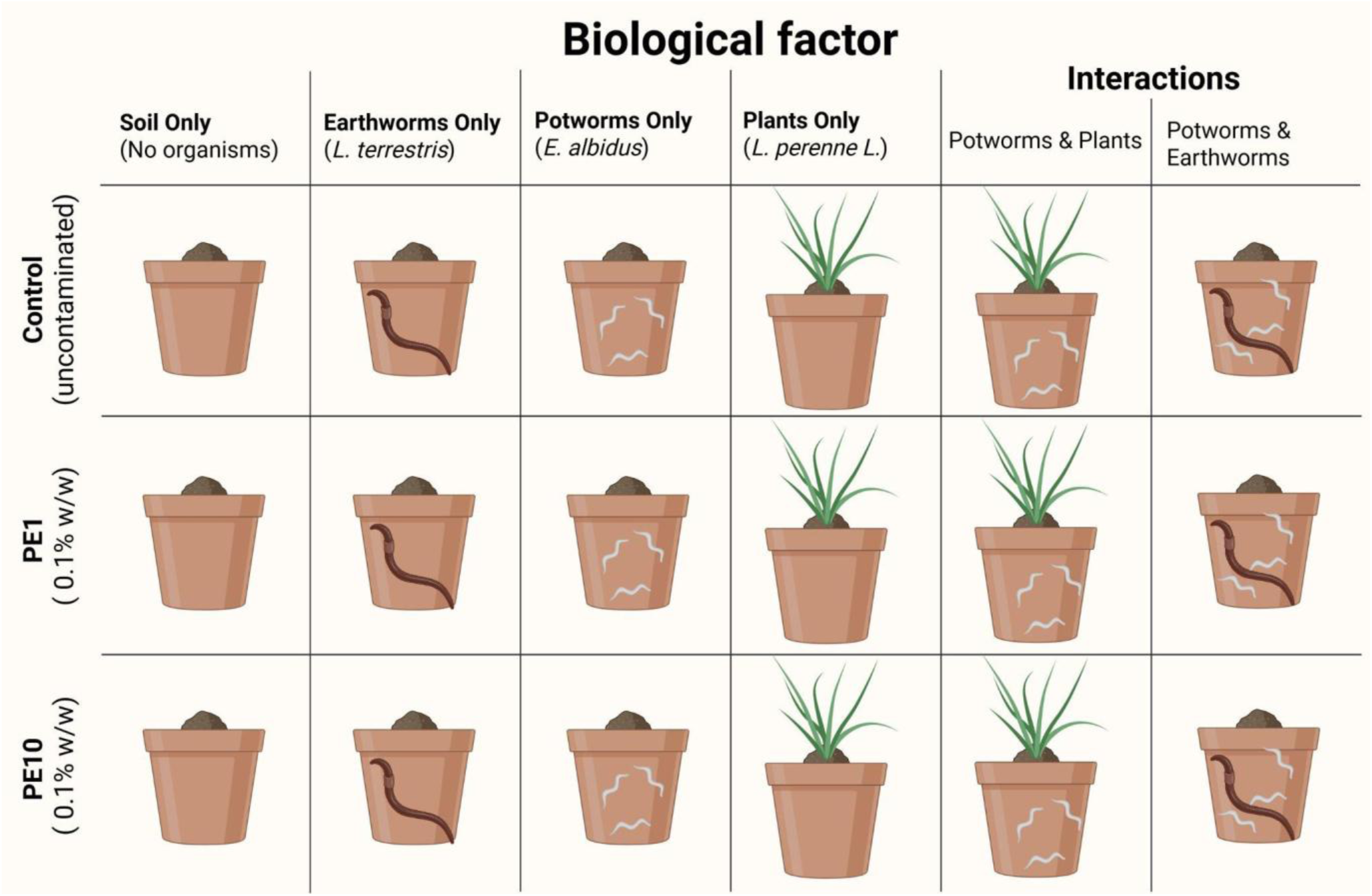
:Graphical representation of the experimental design. For each experimental modality, a total of 4 replicates were analyzed.

**Table 1.** : Effect of additives content in LDPE (PE1/PE10) on soil parameters. Values are mean of 4 replicates ± standard deviation. Asterisks show significant difference compared to control (* : p<0.05, ** : p<0.01 ; statistical tests used are presented in 2.10)

| Parameter | Control | PE1 | PE10 |
| --- | --- | --- | --- |
| **pH** | 6.84 ± 0.07 | 6.94 ± 0.15 | 6.94 ± 0.10 |
| **Conductivity** (μS) | 83.3 ± 3.6 | 82.4 ± 6.8 | 92.4 ± 13.7 |
| **Total carbon** (%) | 0.96 ± 0.00 | **1.07 ± 0.02\*** | 1.04 ± 0.03 |
| **Labile Carbon** (mg C.kg<sup>-1</sup> soil) | 345 ± 19 | 351 ± 4 | 351 ± 5 |
| **SIR** (μg.C-CO<sub>2</sub>/h/g dry soil) | 7.3 ± 3.0 | 7.3 ± 0.7 | 6.3 ± 2.0 |
| **Total Nitrogen** (%) | 0.102 ± 0.002 | 0.104 ± 0.002 | 0.105 ± 0.002 |
| **NO<sub>3</sub><sup>-</sup>** (μg.N-NO<sub>3</sub><sup>-</sup>/g dry soil) | 6.06 ± 1.78 | 5.32 ± 1.45 | 5.71 ± 0.97 |
| **NO<sub>2</sub><sup>-</sup>** (μg.N-NO<sub>2</sub><sup>-</sup>/g dry soil) | 0.029 ± 0.007 | 0.041 ± 0.012 | **0.050 ± 0.005\*** |
| **NH<sub>4</sub><sup>+</sup>** (μg.N-NH<sub>4</sub><sup>+</sup>/g dry soil) | 0.232 ± 0.054 | **0.403 ± 0.102 \*** | **0.484 ± 0.048\*\*** |
| **DEA** (μg.N-N<sub>2</sub>O/h/g/dry soil) | 0.84 ± 0.02 | **1.33 ± 0.18\*** | **1.18 ± 0.16\*** |
| **NEA** (μg.N-NO<sub>2</sub><sup>-</sup> + N-NO<sub>3</sub><sup>-</sup>/h/g dry soil) | 0.574 ± 0.023 | 0.618 ± 0.043 | 0.660 ± 0.088 |

### 2.6 Analyses on MPs characteristics and soil parameters

Samples were analysed using an elemental analyser (Vario Isotop select from Elementar, Lyon, France) to determine C, N concentrations. Data calibration for C, N concentrations were performed using a tyrosine internal reference standard. Analytical accuracy of the C and N elemental analyser was characterised by a relative standard deviation lower than 2%. Soil pH and conductivity were determined according to ISO 103920:1994 (F) and AFNOR X31-113 standards, respectively. Labile carbon was determined according to the method described by Weil et al. (2003) and revised by Gruver (2004)(Weil et al. 2003; Gruver 2004). Soil nitrogen (NO_2_^-^, NO_3_^-^ and NH_4_^+^) concentrations were determined after extraction from soil subsamples (5g soil dry-weight equivalent) with 20 mL of a solution of a 0.01 M calcium chloride (CaCl_2_) solution. The soil was shaken at 140 rpm for 2h at 10 - ◦ C. The soil suspensions were filtered using an Acrodisc Supor®PF with membrane prefilter (0.8/0.2 μm, VWR) and stored at −20°C prior to analysis the NO_2_^−^, NO_3_^−^ and NH_4_^+^ concentrations using colorimetric reaction with sequential analyzer smartchem 200 (Metrohm, France).

### 2.7 Analyses on plant

At the end of the experiment, roots and shoots were separated and weighted using a precision balance. The shoot system was then oven-dried at 65°C for one week to determine the dry biomass.

### 2.8 Analyses on annelid life traits

Surviving *A.caliginosa* were collected and cleaned with water prior to calculating survival rates. Earthworm growth was characterised by measuring fresh earthworm biomass, while reproduction was evaluated by collecting fresh soil at the end of the experiment. The collected soil from each pot was washed through a 1 mm sieve, and the number of eggs per unit of soil was recorded. Half of the collected earthworms were placed on a fast and half were frozen at -80°C. For *E. albidus*, 30 g soil of two subsamples were pooled from each pot, and the number of surviving individuals was counted and weighed. The results were normalized to soil mass to determine organism density. Surviving *E. albidus* were weighted to calculate the average individual biomass.

### 2.9 Analysis of microbial activities

Three microbial activities were assessed in this study: substrate-induced respiration (SIR), denitrification enzyme activity (DEA), and nitrification enzyme activity (NEA). These measurements provide complementary insights into the main soil microbial processes involved in carbon (SIR) and nitrogen (NEA and DEA) cycling. The three activities are briefly described below, while the detailed experimental protocols are provided in Béraud et al. (2025).

#### 2.9.1) Substrate Induced Respiration (SIR)

Substrate-induced respiration (SIR) was measured as CO_2_ production. Carbon dioxide concentrations were measured hourly for 6 h with gas chromatography coupled to a micro-catharometer detector (µGC-R990; SRA Instruments, Marcy L’Etoile, France) and expressed as CO_2_ production (µg C-CO_2_ g^-1^ dry soil h^-1^).

#### 2.9.2) Nitrification Enzyme Activity (NEA)

Potential nitrification enzyme activity (NEA) was measured as NO_2_^-^ and NO_3_^-^ production. The amount of NO_2_^-^ and NO_3_^-^ produced in samples during incubation was measured every two hours for 10 h, for each kinetic sampling point using a Smartchem 200 photometer (AMS Alliance, Villeneuve-la-Garenne, France). The potential nitrification enzyme activity was measured as a NO_2_^-^ + NO_3_^-^ production (µg N-NO_2_^-^ + NO_3_^-^ g^-1^dry soil.h^-1^).

#### 2.9.3) Denitrification Enzyme Activity (DEA)

Denitrification enzyme activity (DEA) was measured as N_2_O production. N_2_O concentrations were measured hourly for 6 h, with a gas chromatograph coupled to a micro-catharometer detector (µGC-R990; SRA Instruments, Marcy L’Etoile, France) and N_2_O produced was expressed in µg N-N_2_O g^-1^ dry soil.h^-1^.

### 2.10 Statistical analysis

Statistical analyses were performed using R (version 4.3.0). Prior to hypothesis testing, all datasets were assessed for normality using the Shapiro–Wilk test and for homogeneity of variances using Levene’s test. Comparisons among multiple groups have been performed :

- comparison between the physico-chemical parameters of uncontaminated soil and those of contaminated soil (PE1 or PE10)
- comparison between the growth of unexposed organisms and the growth of organisms exposed to PE1 and PE10
- comparison between the growth of organisms exposed individually to PE1 and PE10 and the growth of organisms exposed in interaction to PE1 and PE10. Simultaneously, the impact of PE1 and PE10 on growth is tested.

For comparisons among multiple groups, the choice of post hoc procedure was based on data distribution and variance assumptions:

**Tukey’s Honest Significant Difference (HSD) test** was applied following one-way ANOVA when data met assumptions of normality and homogeneity of variances.

When normality was met but variances were unequal, the **Games–Howell test** was employed. For non-normally distributed data, a **Kruskal–Wallis test** was performed followed by **Dunn’s post hoc test with Bonferroni correction** for pairwise comparisons.

The level of statistical significance was set at *p* < 0.05. All data are presented as mean ± standard deviation. Detailed statistical results, including the specific test applied for each comparison, are provided in the figure legends.

Percentages of variations were calculated with the following equation : 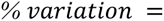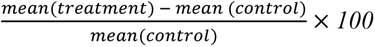. Graphical representations display the average percentage variation in the biomass of organisms in contaminated soils compared to uncontaminated soils (controls), while the graphical representations in **Figure 2** display the average percentage variation in the biomass of organisms in interactions compared to the biomass of organisms without interactions. The error bars represent the standard deviation of the percentage variation. Statistical tests were performed on the untransformed raw data and the results are reported on the ’percentage of variation’ graphics.

**Figure 2.**
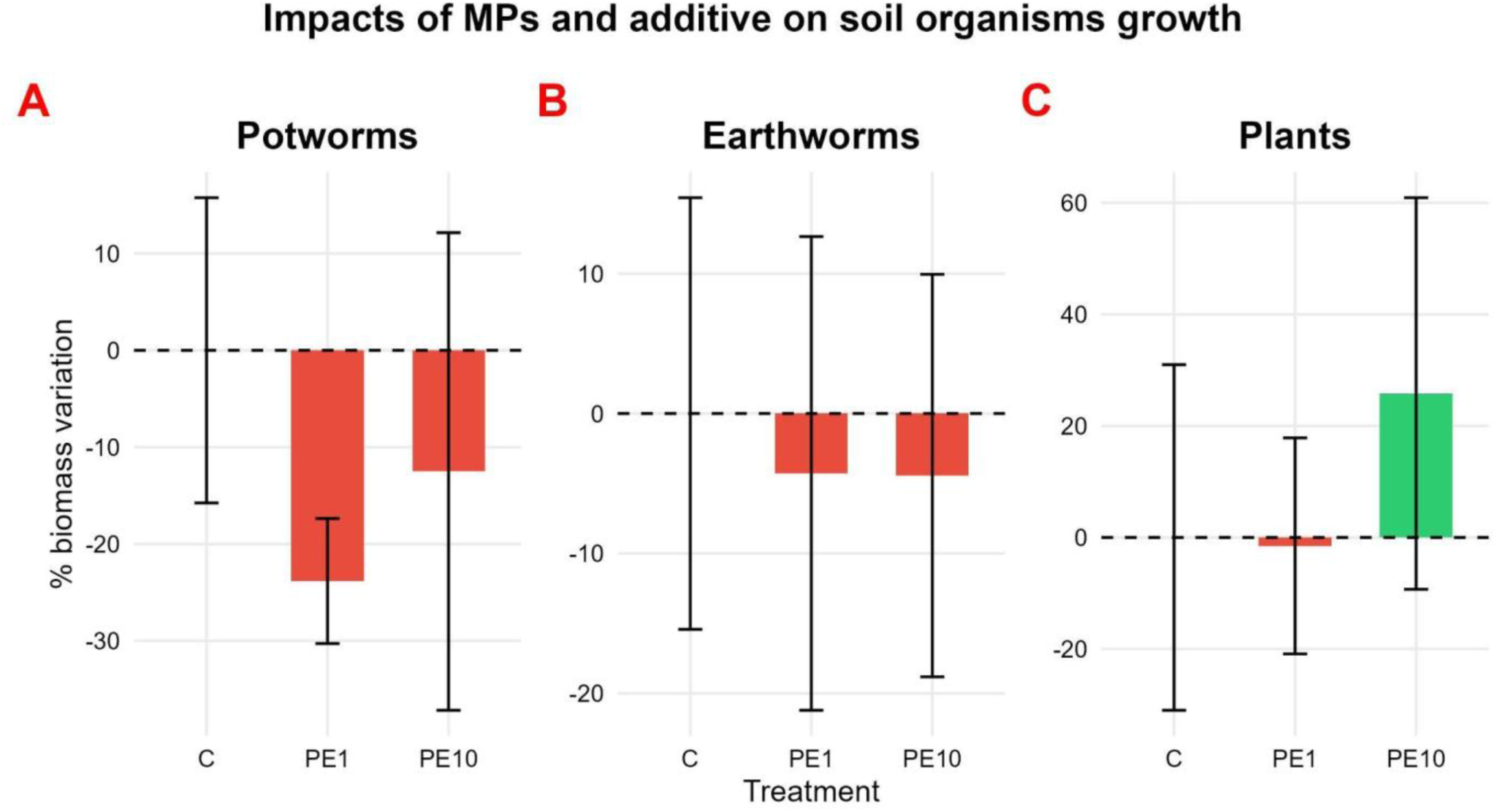
: Impacts of MPs with two different doses of additives (PE1 and PE10) on A : potworm growth, B : earthworm growth, C : plant growth (shoot dry biomass). Letters show significant differences between treatments (p < 0.05) with tests performed on raw data.

## 3. Results

### 3.1 Effects of MPs and additives on soil parameters

Control soil parameters are reported on **Table 2**. No effects of MPs contamination were found on soil pH, conductivity, labile carbon C, total nitrogen N or nitrate concentration (NO_3_ ^-^). However, total C content, nitrites (NO_2_ ^-^) and ammonium (NH_4_ ^+^) concentrations are modified in contaminated soils (**Table 2**).

**Table 2.**
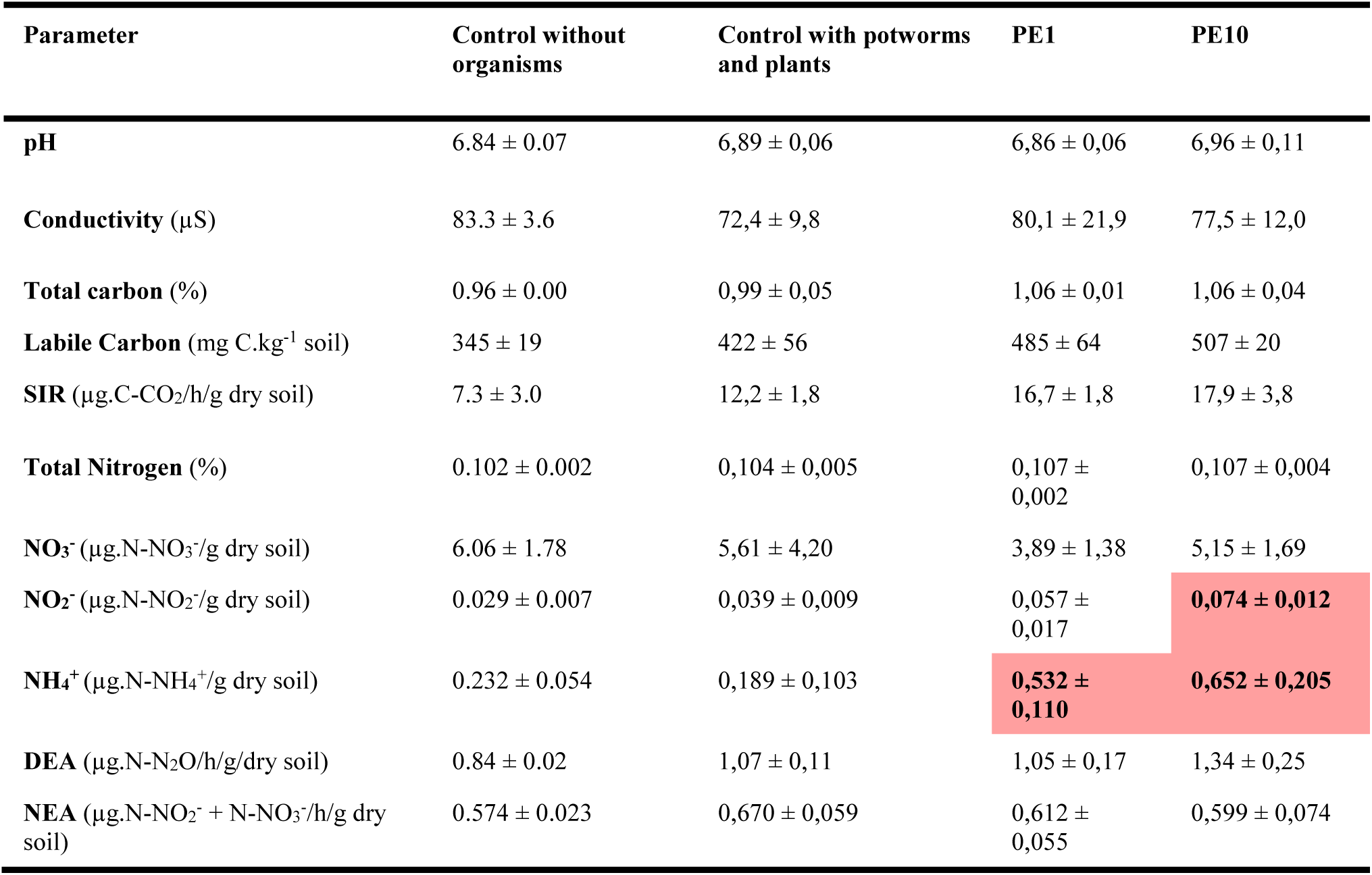
: Effects of MPs on soil parameters in planted soil in presence of potworms. Values are mean of 4 replicates ± standard deviation. Red-colored cells show significant increase compared to control and blue-colored cells show significant decrease compared to control (p < 0.05 ; statistical tests used are presented in 2.10)

| Parameter | Control without organisms | Control with potworms and plants | PE1 | PE10 |
| --- | --- | --- | --- | --- |
| **pH** | 6.84 ± 0.07 | 6,89 ± 0,06 | 6,86 ± 0,06 | 6,96 ± 0,11 |
| **Conductivity** (μS) | 83.3 ± 3.6 | 72,4 ± 9,8 | 80,1 ± 21,9 | 77,5 ± 12,0 |
| **Total carbon** (%) | 0.96 ± 0.00 | 0,99 ± 0,05 | 1,06 ± 0,01 | 1,06 ± 0,04 |
| **Labile Carbon** (mg C.kg<sup>-1</sup> soil) | 345 ± 19 | 422 ± 56 | 485 ± 64 | 507 ± 20 |
| **SIR** (μg.C-CO<sub>2</sub>/h/g dry soil) | 7.3 ± 3.0 | 12,2 ± 1,8 | 16,7 ± 1,8 | 17,9 ± 3,8 |
| **Total Nitrogen** (%) | 0.102 ± 0.002 | 0,104 ± 0,005 | 0,107 ± 0,002 | 0,107 ± 0,004 |
| **NO<sub>3</sub><sup>-</sup>** (μg.N-NO<sub>3</sub><sup>-</sup>/g dry soil) | 6.06 ± 1.78 | 5,61 ± 4,20 | 3,89 ± 1,38 | 5,15 ± 1,69 |
| **NO<sub>2</sub><sup>-</sup>** (μg.N-NO<sub>2</sub><sup>-</sup>/g dry soil) | 0.029 ± 0.007 | 0,039 ± 0,009 | 0,057 ± 0,017 | **0,074 ± 0,012** |
| **NH<sub>4</sub><sup>+</sup>** (μg.N-NH<sub>4</sub><sup>+</sup>/g dry soil) | 0.232 ± 0.054 | 0,189 ± 0,103 | **0,532 ± 0,110** | **0,652 ± 0,205** |
| **DEA** (μg.N-N<sub>2</sub>O/h/g dry soil) | 0.84 ± 0.02 | 1,07 ± 0,11 | 1,05 ± 0,17 | 1,34 ± 0,25 |
| **NEA** (μg.N-NO<sub>2</sub><sup>-</sup> + N-NO<sub>3</sub><sup>-</sup>/h/g dry soil) | 0.574 ± 0.023 | 0,670 ± 0,059 | 0,612 ± 0,055 | 0,599 ± 0,074 |

PE1 and PE10 treatment increased the total C content, respectively by 11 and 8 %, although this was not statistically significant (p>0.05). NH_4_ ^+^ concentrations also increased in both PE1 and PE10 conditions, with the increase being significantly higher in PE10 than in PE1. PE1 treatment significantly increased nitrites concentration by 41%

Soil microbial activities showed no significant effects of MPs contamination on SIR and NEA. However, PE1 and PE10 induced a significant increase of DEA, respectively by 58 and 40 % (**Table 2**).

### 3.2 Impacts of MPs and additives on soil organisms growth

No significant effects of MPs or additives were observed on the growth of soil fauna (potworms, earthworms) or plants when exposed independently to the MPs and the additives (**Fig. 2A, B and C**). For potworms, a non-significant trend toward reduced biomass was observed in the PE1 treatment likely due to the high variability in the control treatment (**Fig. 2A**).

### 3.3 Impacts of MPS and additives on soil parameters and organism growth when they are simultaneously exposed

Potworms showed a significant decrease in average individual biomass of 49 ± 4.1 % when co-exposed with plants (**Fig. 3A**).

**Figure 3.**
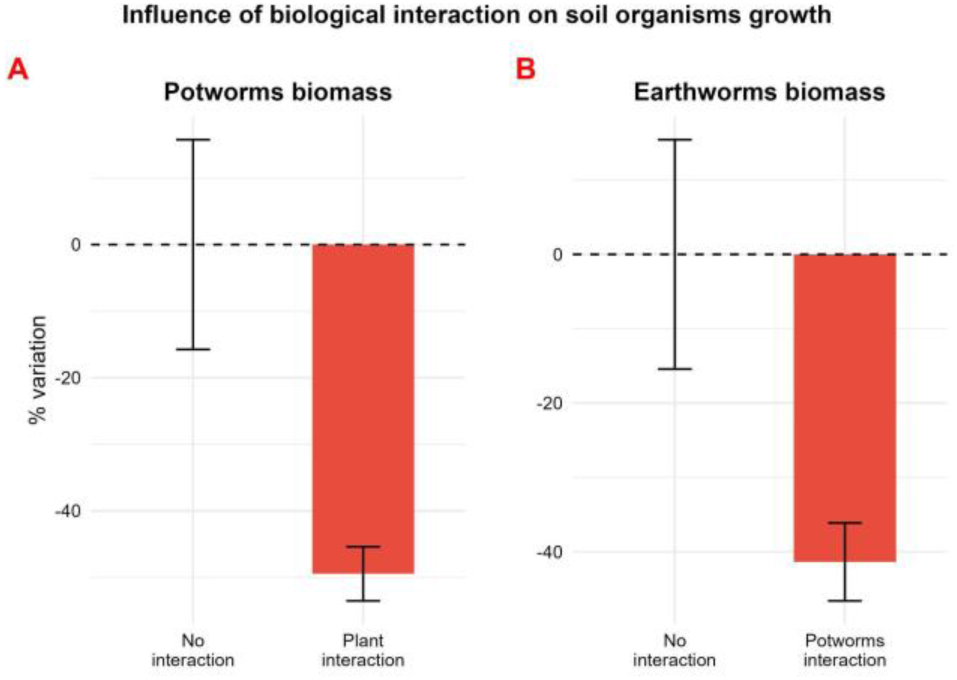
: Graphical representation of influence of biological interaction on soil fauna growth. Bars represents percentage of variation compared to control (A : potworms without interaction, B : earthworms without interaction). Error bars represents standard deviation (SD).

Earthworms exhibited a significant decrease in average individual biomass of 41 ± 5.2% when co-exposed with potworms (**Fig. 3B**). The interaction between potworms and earthworms in uncontaminated soil also enhanced SIR by 71%. Similarly, NO_3_ ^-^ concentrations increased by 175% when earthworms and potworms are incorporated in uncontaminated conditions. However, these increases were not statistically significant due to the high variability in the interaction control group (SD = 8.72 µg N- NO_3_ ^-^ g^-1^dry soil).

No significant effects of PE1 or PE10 treatments were observed on potworm growth when exposed alone to MPs (**Fig. 1A**). However, the potworm average individual biomass significantly increased when exposed to PE10 with plants (**Fig. 4**, p<0.05). The average individual biomass was 1,16 ± 0,09 mg in Control + plants and 2,01 ± 0,27 mg in the PE10 +plants treatments.

**Figure 4.**
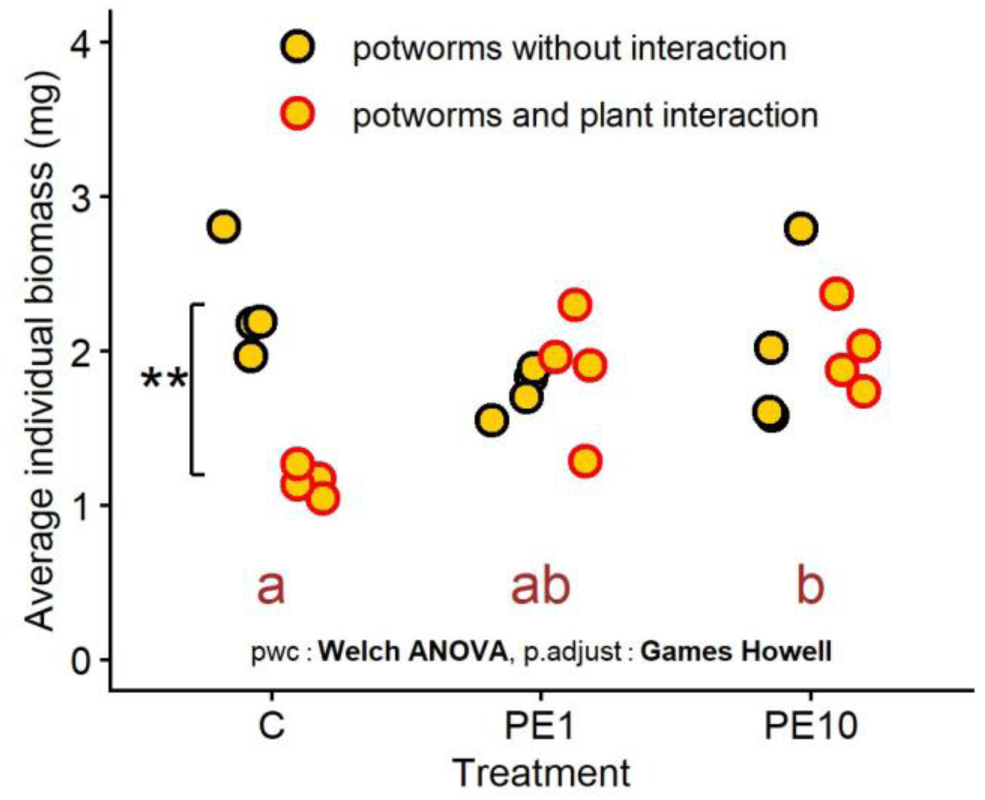
: Potworms growth in unplanted and planted condition. Asterisks show significant difference with the presence of plants compared to soil with potworms alone (* : p<0.05, ** : p<0.01 ; statistical tests used are presented in 2.10). Different letters show statistical differences among the treatments compared to control C uncontaminated).

PE10 treatment increased nitrite concentration by 90 % and ammonium concentration by 243 %. A similar increase of NH ^+^ concentration was also observed in the PE1 treatment (181 %).

No significant effects were observed following PE1 or PE10 treatments on earthworms alone (**Fig. 2B**). However, the earthworm average individual biomass increased when co-exposed to PE10 with potworms (**Fig. 5** p<0.05). The average individual biomass was 284,25 ± 25,4 mg in control+potworms and 369,75 ± 25,8 mg in PE10 +potworm treatment.

**Figure 5.**
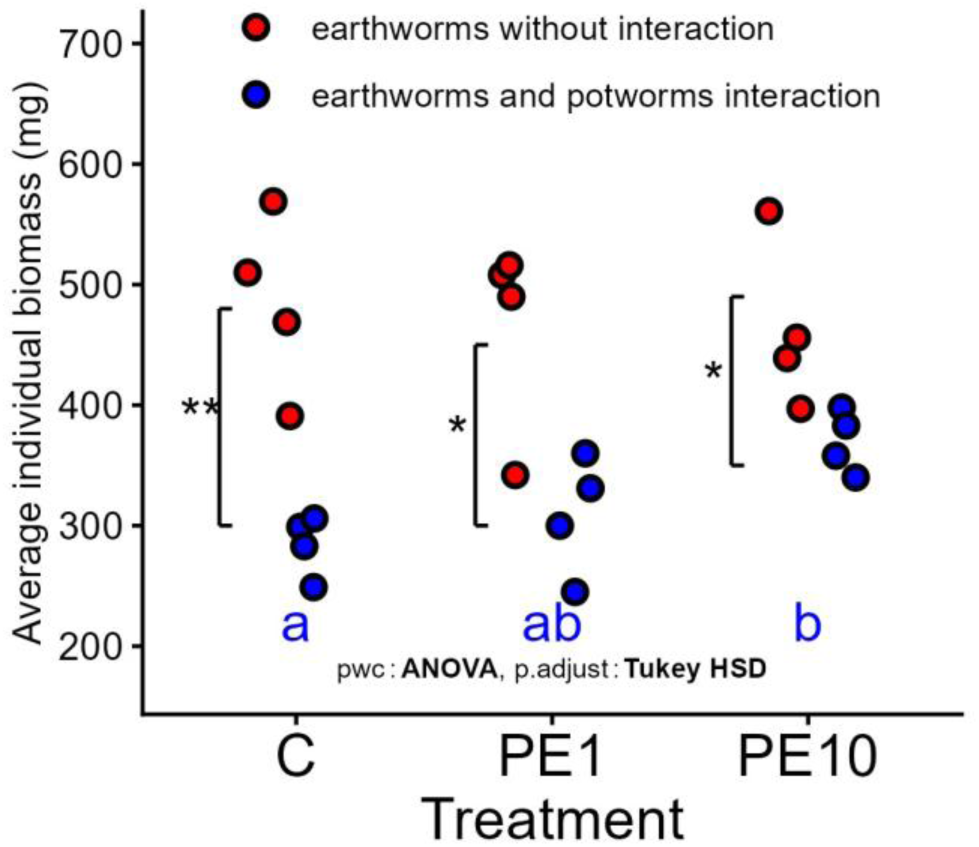
: Earthworm growth with and without potworms. Asterisks show significant difference with the presence of potworms compared to soil with earthworms alone (* : p<0.05, ** : p<0.01 ; statistical tests used are presented in 2.10). Different letters show statistical differences among the treatments compared to control C uncontaminated).

When earthworms and potworms were co-exposed to PE1, soil pH significantly decreased by 4 % and total C content significantly increased by 8%. Addition of PE10 in soil within earthworms and potworms significantly increased total C content (6 %), total N (5 %), NO_3_^-^ concentration (229 %) and NH_4_^+^concentration (173 %) (**Table 3**).

**Table 3.** : Effects of MPs on soil parameters in unplanted soil in presence of earthworms and potworms. Values are mean of 4 replicates ± standard deviation. Asterisks show significant differences in the uncontaminated soil with interaction compared to the uncontaminated soil without interaction (p<0.05). Red-colored cells show significant increase in contaminated soil with interaction compared to control with interaction and blue-colored cells show significant decrease compared to control with interaction (p < 0.05 ; statistical tests used are presented in 2.10)

| Parameter | Control without organisms | Control with earthworms and potworms | PE1 | PE10 |
| --- | --- | --- | --- | --- |
| pH | 6.84 ± 0.07 | 6,70 ± 0,08 | 6,45 ± 0,06 | 6,54 ± 0,11 |
| Conductivity (μS) | 83.3 ± 3.6 | 103,0 ± 26,0 | 129,78 ± 24,3 | 140,8 ± 17,2 |
| Total carbon (%) | 0.96 ± 0.00 | 0,96 ± 0,03 | 1,04 ± 0,02 | 1,02 ± 0,03 |
| Labile Carbon (mg C.kg<sup>-1</sup> soil) | 345 ± 19 | 422 ± 168 | 439 ± 23 | 514 ± 62 |
| SIR (μg.C-CO<sub>2</sub>/h/g dry soil) | 7.3 ± 3.0 | 12,5 ± 0,8* | 13,3 ± 4,5 | 12,9 ± 0,4 |
| Total Nitrogen (%) | 0.102 ± 0.002 | 0,102 ± 0,002 | 0,106 ± 0,003 | 0,107 ± 0,002 |
| NO<sub>3</sub><sup>-</sup> (μg.N-NO<sub>3</sub><sup>-</sup>/g dry soil) | 6.06 ± 1.78 | 16,64 ± 8,72 | 27,84 ± 10,10 | 54,73 ± 17,20 |
| NO<sub>2</sub><sup>-</sup> (μg.N-NO<sub>2</sub><sup>-</sup>/g dry soil) | 0.029 ± 0.007 | 0,030 ± 0,002 | 0,031 ± 0,004 | 0,033 ± 0,007 |
| NH<sub>4</sub><sup>+</sup> (μg.N-NH<sub>4</sub><sup>+</sup>/g dry soil) | 0.232 ± 0.054 | 0,247 ± 0,047 | 0,402 ± 0,071 | 0,675 ± 0,223 |
| DEA (μg.N-N<sub>2</sub>O/h/g/dry soil) | 0.84 ± 0.02 | 0,74 ± 0,22 | 0,89 ± 0,43 | 0,53 ± 0,11 |
| NEA (μg.N-NO<sub>2</sub><sup>-</sup> + N-NO<sub>3</sub><sup>-</sup>/h/g dry soil) | 0.574 ± 0.023 | 0,628 ± 0,053 | 0,833 ± 0,259 | 1,136 ± 0,060 |

## 4. Discussion

### 4.1 Effects of MPs and additives on soil physico-chemical properties

Slight increases in soil pH were observed in the contaminated soil; however, the variability among replicates prevented these differences from being statistically significant. MPs have been reported to influence soil pH, either by increasing (Qi et al. 2020) it or decreasing it (Palansooriya et al. 2022). Moreover, Yue et al. (2023) suggested that both the direction and magnitude of MP effects on soil pH may depend on exposure time and polymer type.

We observed that the incorporation of MPs in the soil significantly influenced several soil parameters including total carbon, denitrification (DEA), and nitrites and ammonium (NH_4_^+^) content. The PE1 and PE10 treatments significantly stimulated DEA, although there were no observed changes in soil nitrate concentration. The increase of total carbon is most likely attributable to the presence of the MPs, which represented an increase of 0.11 and 0.08% respectively. This rise is probably linked to the addition of 0.1% of MPs, mainly composed of carbon (Rillig 2018). However, there is no reliable analytical method to distinguish soil organic carbon from the carbon derived from MPs (Rillig et al. 2021). Interestingly, the lower total carbon content observed in the PE10 treatment compared to PE1 could reflect variations in polymer formulation, particularly a higher proportion of additives containing carbon, as well as other elements such as nitrogen, oxygen, phosphorus and hydrogen (Appendix 2). Regarding NH_4_^+^ soil content, the incorporation of MPs induced a significant increase, consistent with previous findings (J. Zhang et al. 2022). The mechanism underlying this increase remains unclear. Plastic residues may act as a physical barrier limiting root penetration and thereby reducing plant nitrogen uptake, ultimately leading to NH_4_^+^accumulation (Zumilaiti et al. 2017). Other studies have also reported an increase in soil water-holding capacity in soils contaminated with mulch film residues. Enhanced water retention can stimulate soil enzyme activities, especially those involved in ammonification (F. Ma et al. 2015), which could explain the higher ammonium levels observed (D. Zhang et al. 2017). Moreover, the incorporation of PE film has been shown to alter the soil microbial community structure, notably by decreasing the abundance of ammonia-oxidising bacteria (AOB), which are responsible for converting ammonium into nitrate. Reduced AOB abundance and activity may limit ammonium oxidation and promote its accumulation in the soil (Gao et al. 2021). Li et al. (2025) further suggested that MPs can modulate enzyme activity by binding to or displacing metal ions associated with their functional groups (thiol, carboxyl, imidazole, etc.), resulting in complex enzymatic response to MP exposure. In our experiment, we observed an accumulation of NH_4_^+^ in contaminated soil, but no significant effect on nitrification (NEA) was detected (Tab 2). However, these mechanisms remain largely hypothetical and have not yet been experimentally validated. In our experiment, we observed an accumulation of NH_4_^+^ in contaminated soil, but no significant effect on NEA was detected (Tab 2). By contrast, the PE1 and PE10 treatments significantly stimulated DEA, although no changes in soil nitrate concentration were observed.

### 4.2 Interspecific interactions under uncontaminated conditions

When both earthworms and potworms are present under uncontaminated conditions, the average individual biomass of earthworms decreased in the presence of potworms (**Fig 3B**). Our results are consistent with the literature, which also reported a reduction in earthworm biomass when co-incubated with potworms (Sandor et Schrader 2012). It should be noted that laboratory experiments on annelid competition often use organism densities higher than those found in natural conditions, which can lead to reduced individual biomass due to increased competition for resources (Eriksen-Hamel et Whalen 2007). We also observed a significant increase in substrate induced respiration (SIR) when both earthworms and potworms were present compared to the control condition. Similar patterns have been reported previously, with enhanced soil respiration following earthworm addition, largely attributed to stimulation of microbial activity (Bohlen et Edwards 1995). When coexisting, potworms and earthworms therefore appear to stimulate nitrogen mineralization. However, their interaction seems to modify nitrogen cycling dynamics. Indeed, no decrease of nitrite concentrations was observed when the organisms were present separately. Since potworms are known to graze on microorganisms (Hedlund et Augustsson 1995), the presence of earthworms may have altered their feeding behavior. This could have reduced *Nitrobacter* populations, thereby limiting nitrification and resulting in reduced N mineralization and slight nitrite accumulation. This interpretation aligns with their known feeding preferences: earthworms are approximately 80% saprovorous and 20% microbivorous, whereas potworms are about 80% microbivorous and 20% saprovorous (Didden 1993).

When both plants and potworms are present under uncontaminated conditions, we observed a negative effect of plants on potworm biomass (**Fig 3A**). While numerous studies have explored the effects of earthworms on plant productivity, the reciprocal influence of plants on annelids growth remains poorly documented. Potworms, particularly enchytraeids, are known to play an important role in organic matter decomposition, soil structuring, and aeration (Didden 1993). These activities are usually beneficial to plant growth (Cole et al. 2002). It is commonly acknowledged that plants stimulate microbial activity through root exudates (Quigley et al. 2025), enhancing heterotrophic microbial communities. In our study, we found that plants negatively affected potworms biomass, apparently contradicting these reports. As potworms are primarily microbivorous, plant presence should theoretically promote potworm growth by providing additional microbial resources through the rhizosphere stimulation. Based on our experimental design, we hypothesise that the root systems enhance the spatial constraints within microcosm, acting as a physical barrier that restricts potworms movement and habitat, thereby explaining the reduction in average individual biomass.

### 4.3 Modulation of interspecific interactions by MPs and additives

Individual exposure to MPs, regardless of additives concentration, had no impact on the growth of soil organisms compared to the non-contaminated soil. This has been reported for potworms exposed from 2 to 5% of MPs (Šmídová et al. 2024), earthworms (Mohammed et al. 2025; Rodriguez-Seijo et al. 2017), and plants (no effect on *Lolium multiflorum* with HDPE (Esterhuizen et al. 2022)). Although there is no data available for an LDPE exposure to *Lolium perenne* L., Fajardo et al. (2022) reported growth inhibition in maize following 0.1% PE exposure. Significant differences emerged during interspecific interactions. The strong negative effects of plants on potworms (section 4.2) and potworms on earthworms) in uncontaminated soil were abolished under PE1/PE10 treatments. Specifically, potworm biomass increased significantly under PE10 + plants, and earthworm growth rose under PE10 + potworms. No significant differences occurred between control and PE1 treatments, suggesting additive concentration, rather than MPs themselves, drives these effects. These results suggest that MPs, regardless of additive concentration, may mitigate the adverse effects of plants on potworm growth. As no significant differences were found between the control and PE1 treatment, it is reasonable to hypothesize that the observed mitigation is primarily linked to the additives’ composition rather than to the MPs themselves. Potworms exerted a significant negative effect on earthworm growth in uncontaminated conditions (**Fig. 5**). However, when both species were co-exposed to PE10, earthworm growth significantly increased relative to uncontaminated conditions. These findings suggest that high additives’ concentration may alleviate the negative influence of potworms on earthworm growth. Again, as no significant difference was observed between the control and PE1 treatments, these effects are likely driven by the additives’ concentration rather than by the presence of MPs themselves.

Under the PE10 exposure with both plants and potworms, Total Carbon, Total nitrogen, soil nitrate and ammonium concentrations increased compared to uncontaminated conditions. These results indicate that the reduction of the potworm-induced inhibition of earthworm growth following PE10 exposure is closely related to altered soil chemical properties, particularly nitrogen dynamics. However, the nitrate accumulation pattern differed from that observed in the earthworm-potworm interaction without plants. Although *L. perenne* is known to preferentially uptake ammonium over nitrate (Clarkson et al. 1986; Spratt et Gasser 1970), no significant effect of plants was observed on soil ammonium concentrations. The lower soil nitrate concentrations observed in this case, suggest a preferential uptake of nitrate, likely due to the lower ammonium availability in the studied soil. Nitrite and ammonium soil concentrations significantly increased under PE10 exposure with earthworms and potworms, compared to the control. This pattern indicates that the mitigation of plant-related negative effects on potworms in this condition is probably associated with changes in soil properties, notably related to nitrogen content.

Recent studies show that plastic additives can leach and accumulate in soils (Xu et al. 2024). In our study, LDPE- associated additives increased soil ammonium, unlike Zhou et al. (2024), who reported higher total nitrogen but lower soil ammonium levels. This discrepancy is likely related to differences in additives’ composition, as our formulation included nitrogen-containing additives such as Chimassorb 944, Erucamide and Oleamide. Although little is known about their specific soil effects, Erucamide and Oleamide have been linked to enhanced denitrification activity (Lu et al. 2014; Sun et al. 2016). The effects of Oleamide, however, remain largely unexplored (Jug et al. 2020; Cooper et Tice 1995). Oleamide is a naturally occurring endogenous lipid in mammals, structurally related to endocannabinoids (Hiley et Hoi 2007). Luo et al. (2025) recently reported that Oleamide produced by *Streptomyces lydicus* JCK-6019 in the cucumber rhizosphere enhanced plant resistance against two pathogenic fungi. The leaching of such additives may therefore explain the observed stimulation of denitrifying activity (DEA) in contaminated soils (Table 1). This result suggests that these compounds could influence the nitrogen cycle. Furthermore, the potential degradation and/or transformation of these compounds might release additional mineral nitrogen. In contrast, currently there is no data available on the soil impacts of Chimassorb 944, despite evidence that it can leach and degrade in the environment (Haider et Karlsson 1999). The findings of this study demonstrate that when evaluating the toxicity of MPs to the soil environment, consideration must be given to the additives’effects on soil organisms. Yet, few information is available regarding the individual ecotoxicity of our additives on soil organisms. The only ecotoxicological data is mostly extracted from registration documents (data are presented in S4). In order to quantify the influence of additives on MPs toxicity, research on the individual contribution of each used additive is needed.

## 5. Conclusion

In this study, individual growth of earthworms, potworms and *Lolium perenne* remained unaffected by PE1 or PE10 microplastic exposure. However, significant biotic interactions emerged under uncontaminated conditions: plants significantly reduced potworm growth, and potworms reduced earthworm biomass. Remarkably, PE10 contamination alleviated these antagonistic effects, concurrently with disruption of soil N cycling (elevated soil ammonium concentration and stimulated DEA). PE1 triggered weaker responses than PE10 exposure, implicating that additives leaching over MP particles themselves. This underscores the critical need for multi-species designs to detect indirect MP effects on soil functioning, as single-species studies consistently report negligible direct toxicity. Therefore, future research must quantify the additives’ degradation pathways and their mineral N release, in addition to assessing the chronic multi-trophic effects under realistic field-relevant concentrations, and investigate polymer-specific additive interactions, as formulation governs toxicity. Multi-species microcosms reveal hidden MP risks that single-species toxicology overlooks, particularly when shifts in additive-driven soil chemistry mediate ecological responses.

## Credit authorship contribution statement

**Blanchard Adrien:** Conceptualization, Methodology, Investigation, Writing - original draft, **Faburé Juliette:** Supervision, Writing - review & editing, Validation, Funding acquisition, **Bamière Antoine:** Methodology, Resources, **Breuil Sébastien**: Methodology, Resources, **Creuzé des Châtelliers Charline**: Methodology, Resources, **Delort Abigaïl:** Methodology, Resources, **Etiévant Véronique**: Methodology, Resources, **Gervaix Jonathan:** Methodology, Resources, Validation, **Marret Manon**: Methodology, Resources, **Plessis Cédric**: Methodology, Resources, **Cantarel Amélie A.M.:** Supervision, Writing - review & editing, Validation, Funding acquisition, **Richaume Agnès**: Supervision, Writing - review & editing, Validation, Funding acquisition.

## Data availability

Data will be made available on request.

## Acknowledgments

Authors acknowledge financing from ANR (e-DIP, ANR-21-CE34-0017) and INRAE Biosefair Metaprogram. The QUALIAGRO field experiment is part of the SOERE-PRO (network of long-term experiments dedicated to the study of impacts of organic waste product recycling) integrated as a service of the “Investment in the Future” infrastructure AnaEE-France, overseen by the French National Research Agency (ANR-11-INBS0001). The field experiment is financially supported by Veolia. Authors would like to thank Anaee, Veolia and the SOERE-PRO for the access to the stored soil and compost samples of the long-term field experiment, especially Camille Resseguier and Grigorios Andronidis. Authors would like to thank Microbial Activities in the Environment (AME) platform and Environmental Genomic Platform (PGE) at the LEM (Lyon, France), for the soil microbial measurements. Authors would like to thank Patrick Navard and Veronika Khodyrieva for the production of plastic material.

## Abbreviations

MP(s): Microplastic(s)
LDPE: Low-Density Polyethylene
PE: Polyethylene
PE1: LDPE MP with standard additives dose
PE10: LDPE MP with 10 time additives dose
P: Plant
TC: Total Carbon
SIR: Soil Induced Respiration
TN: Total Nitrogen
DEA: Denitrification Enzyme Activity
NEA: Nitrification Enzyme Activity

