## Supplementary material for "Ecological Impacts of Additive-Enriched LDPE Microplastics in Agricultural Soils: Single and Multi-Species Assessments"

S1 : global primary plastic production by polymer in 2019 (https://ourworldindata.org/grapher/plastic-production-polymer)

| Plastics polymer | amount used (Mt) | |
|---|---|---|
| Other | 80,96 | 17,60972 |
| Marine coatings | 0,541 | 0,117674 |
| LDPE, LLDPE | 54,303 | 11,81152 |
| HDPE | 55,544 | 12,08145 |
| PP | 72,805 | 15,83592 |
| PS | 21,116 | 4,592971 |
| PVC | 51,392 | 11,17835 |
| PET | 24,918 | 5,419949 |
| PUR | 18,032 | 3,922166 |
| Fibres | 60,448 | 13,14813 |
| Road marking coatings | 0,682 | 0,148343 |
| Elastomers (tyres) | 7,734 | 1,682233 |
| Bioplastics | 2,326 | 0,505932 |
| ABS, ASA, SAN | 8,944 | 1,945422 |
| Total | 459,746 | 99,99978 |

S2 : Additive composition of tested plastic material

| Additives | PE1 | PE10 | Chemical formula | Function | CAS | Chemical structure |
|---|---|---|---|---|---|---|
| Irganox 1076 | 0,2% | 2% | $C_{35}H_{62}O_3$ | Antioxidant stabilizer | 2082-79-3 | 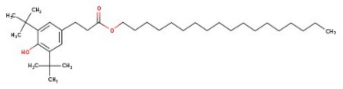 |
| Irgafos 168 | 0,1% | 1% | $C_{42}H_{63}O_3P$ | Secondary antioxidant | 31570-04-4 | 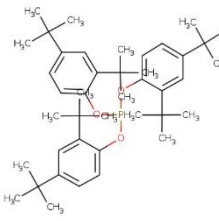 |
| Chimassorb 944 | 0,5% | 5% | $C_{35}H_{68}N_8$ | Light stabilizer | 71878-19-8 | 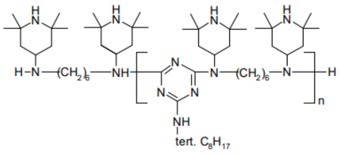 |
| Erucamide | 0,05% | 0,5% | $C_{22}H_{43}NO$ | Slip agent | 112-84-5 | 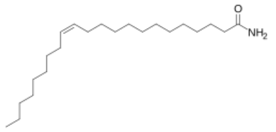 |
| Oleamide | 0,05% | 0,5% | $C_{18}H_{35}NO$ | Slip agent | 301-02-0 | 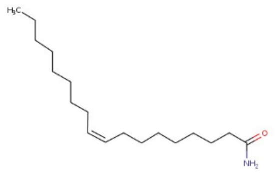 |
| GMS | 0,25% | 2,5% | $C_{21}H_{42}O_4$ | Lubricant, anti-static, plasticizer, anti-aging | 31566-31-1 | 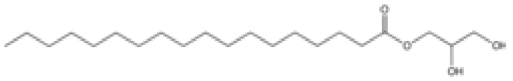 |

S3 : Soil physicochemical properties after sampling at the parcel

| Parameter | Unit | Value (± SD) |
|---|---|---|
| pH | - | 6.6 ± 0.1 |
| Corg | % | 1.03 ± 0.03 |
| Ntot | % | 0.1 ± 0.01% |
| CEC | $cmol^{+}/kg$ | 9.3 ± 0.7 |
| WRC | % | 27.83 |
| Sand [50-2000µm] | % | 6.7 |
| Silt [2-50 µm] | % | 78.15 |
| Clay [< 2µm] | % | 15.15 |

S4 : Available soil ecotoxicological datas for additives used in PE1 and PE10

| **Additives** | Data type | Organism | Value | Source |
|---|---|---|---|---|
| Irganox 1076 | CL/CE50 | Plant | 24 mg/kg | UNEP p20 |
| Irgafos 168 | NOEC | Earthworms | 1000 mg/kg | ECHA |
| Chimassorb 944 | No data available | | | |
| Erucamide | NOEC | Fish | 105 µg/L | ECHA |
| Oleamide | EC50 | cyanobacteria | 8.60 mg/L | Shao et al, 2015 |
| GMS | NOEC | Earthworms | >1000 mmg/kg | ECHA |
